# Batch Effect Correction for Neuroimaging Data with Heterogeneous Spatial Correlations

**DOI:** 10.64898/2026.06.03.729396

**Authors:** Ryan Xie, Dhivya Srinivasan, Gareth A. Harman, Christos Davatzikos, Russell T. Shinohara, Haochang Shou

## Abstract

Magnetic resonance imaging scans are effective tools for unveiling brain structures and understanding pathology for complex neurodevelopmental and aging processes. The spatial correlations among various brain regions reveal critical information and insights into the mechanisms of brain functions and their associations with cognitive abilities. Large-scale neuroimaging studies that acquire or aggregate imaging scans from multiple sites have become increasingly popular. Doing so enhances the diversity of study samples and robustness of study findings, and increases the statistical power of any analysis conducted for the biological hypotheses of interest. However, collecting images across different sites introduces non-biological variability attributed to differences in imaging protocol and configurations, known as batch effects. While there are methods to perform this batch effect correction, there are limited methods that directly account for the spatial patterns found in images of the brain. We develop Covariance-Aware Multivariate (CAM) ComBat that accounts for such spatial correlation across high-dimensional features, which could be heterogeneous across batches. We also propose a computationally efficient alternative of CAM-ComBat, Spatially-Informed Iterative Block (SIB) ComBat, that is scalable for very high dimension of features. We show that these methods outperform existing batch effect correction methods through simulation studies and an application to real neuroimaging data.

## 1 Introduction

Magnetic resonance imaging (MRI) has proven to be a valuable technology in understanding brain pathology and development. The spatial patterns in these images inform the mechanisms and processes underlying brain functions and diseases-related changes [Botz et al., 2023, McIntosh et al., 1996]. Voxel-based morphometry has been especially useful for its ability to detect structural abnormalities and clinically relevant changes in gray and white matter [Mechelli et al., 2005]. Many studies have leveraged voxel-based morphometric analysis to associate changes in the brain with neurological and psychiatric conditions such as schizophrenia, Parkinson’s disease, and epilepsy [Kubicki et al., 2002, Price et al., 2004, Keller et al., 2004]

Pooled analyses by aggregating neuroimaging scans acquired across multiple sites and studies have become an increasingly popular strategy. Doing so not only helps to enhance the diversity of study samples and improve generalizability of the findings, but also boosts the statistical power of any subsequent analysis for the biological hypotheses of interest via larger sample size. Various consortia have implemented this multi-site approach for collecting neuroimaging data [Mueller et al., 2005, Sudlow et al., 2015]. For example, the imaging-based SysTem for AGing and NeurodeGenerative diseases (iSTAGING) consortium collects multimodal MRI scans from over 70,000 individuals across 32 study sites, with the goal of better understanding the mechanism and manifestation of aging and neurodegeneration with respect to the brain [Pomponio et al., 2020, Habes et al., 2021]. However, collecting images at different sites introduces potential non-biological variability attributed to differences in imaging protocol and configurations, known as batch effects [Fortin et al., 2017, 2018]. Previous work has identified inter-scanner variability in voxel-based morphometry and shown it can affect the results of many subsequent analyses [Takao et al., 2011].

Many methods to address these effects aim to remove variability in the means and variances between sites [Fortin et al., 2016, Rao et al., 2017, Yamashita et al., 2019]. A state-of-the-art imaging harmonization method is ComBat, initially formulated to eliminate batch effects for microarray gene expression data prior to its adaptation to neuroimaging data. ComBat uses an empirical Bayes framework to estimate and then adjust for site-specific effects [Johnson et al., 2007]. ComBat has been successfully applied to harmonize various types of neuroimaging data, including cortical thickness and diffusion tensor imaging data [Fortin et al., 2017, 2018]. In its method, ComBat assumes that features are independent of one another, and therefore, does not directly account for differences in correlation patterns between sites.

Multiple methods have been developed to better address this correlation between features. Chen et al. (2021) demonstrated the presence of site covariance differences in cortical thickness measurements from the Alzheimer’s Disease Neuroimaging Initiative (ADNI), and subsequently developed CovBat to mitigate these effects. It expands upon the existing ComBat framework by assuming correlation between residuals and shifting within-site covariances to the pooled covariance through modification of principal component scores [Chen et al., 2022]. While this method proved effective for harmonizing this data, this assumption about the covariance site effects may be less suitable when data present large covariance heterogeneity between batches, especially when dealing with higher dimensions of data. Larger heterogeneity can occur if the batch-specific covariances cannot be captured within the same principal component space, which CovBat assumes. Figure 12 in the Appendix illustrates this idea, in which the similarity of the first ten principal component eigenvectors were compared between data from two study sites. For this specific example, most of the dot products between eigenvectors were close to zero, demonstrating the batch-specific covariances lie in different principal component spaces. Zhang et al. (2024) proposed spatial autocorrelation normalization (SAN) to harmonize vertex level cortical thickness data using spatial Gaussian processes. However, they acknowledged the practical limitations of applying SAN to data with a larger number of features [Zhang et al., 2024]. In addition, calculating covariances using spatial autocorrelation functions may be less flexible for other types of heterogeneity between batches, especially if such heterogeneity arises from batch-specific covariances lying in different principal component spaces.

We developed Covariance-Aware Multivariate (CAM) ComBat to account for heterogeneous spatial correlations in brain images, and show that it outperforms existing batch effect correction methods through simulation tests. To further allow the algorithm to be scalable with higher dimension of features at the voxel-level, we then propose an extension of the CAM ComBat framework to ease the computational burden. We demonstrate through simulation experiments and an application to voxel-wise gray matter density of RAVENS maps from the iSTAGING consortium at the scale of 640,000 features, that our proposed method achieves superior performance compared to other existing batch effect correction methods. Throughout this manuscript, batch and site effects are terms that will be used interchangeably.

## 2 Methods

We first discuss the general framework of ComBat. In summary, ComBat uses an empirical Bayes framework to estimate and then adjust for mean and variance site-specific effects. Specifically, if *Y*_*ijv*_ is the *v*^*th*^ feature from the *j*^*th*^ individual of the *i*^*th*^ batch, ComBat assumes these features can be represented by the following model

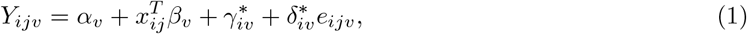

in which *α*_*v*_ is the intercept, *x*_*ij*_ is the vector of covariates that are biologically relevant to the outcome, *β*_*v*_ are the regression coefficients, *γ*_*iv*_ is the additive site effect, *δ*_*iv*_ is the multiplicative site effect, and *e*_*ijv*_ are independent, normally distributed error terms. *α*_*v*_ and *β*_*v*_ are first estimated based on their feature-wise ordinary least squares estimates 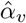 and 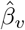 . To estimate *γ*_*iv*_ and *δ*_*iv*_, ComBat assumes independent normal priors on *γ*_*iv*_ and independent inverse gamma priors on *δ*_*iv*_. Using the method of moments to estimate the hyperparameters of these distributions, estimates of the additive and multiplicative site effects, 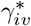and 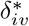, are derived based on the expected value of their respective posterior distribution. The final harmonized dataset 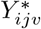 is calculated as follows [Johnson et al., 2007, Fortin et al., 2017, 2018]

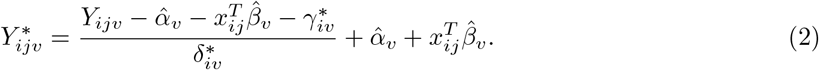

CovBat builds upon this model by assuming the residuals have mean 0 and site-specific covariance Σ_*i*_. To address the covariance site effects, principal component analysis is performed on these full data residuals and the resulting principal component scores are shifted and scaled to get rid of site-specific means and variances in the scores.

In the next section, we expand upon the ComBat framework to specifically address heterogeneous correlations between sites. We introduce two sets of methods: CAM ComBat and Spatially-informed Iterative Block (SIB) ComBat.

### 2.1 CAM ComBat

We assume a similar model as in equation (1) for *Y*_*ijv*_ from batch *i*. However, unlike in conventional ComBat where *e*_*ijv*_ is assumed to follow 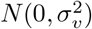, we model site-specific covariance patterns in *e*_*ij*_ and allow them to be correlated following a multivariate normal distribution. That is, 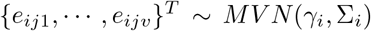. We assume that 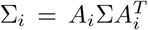 and allow site-specific rotations *A*_*i*_ to be applied onto the common underlying covariance matrix Σ that is shared across sites. This is in contrast to CovBat, in which we assume that Σ_*i*_ can be approximated as ΦΛ_*i*_Φ^*T*^, where Φ are the matrix of the first *q* eigenvectors shared between batches and estimated from the pooled sample covariance, and Λ_*i*_ is a diagonal matrix of eigenvalues for batch *i* and are correspond to the variances of the principal component scores. Compared to CovBat, CAM ComBat’s assumption enables covariances to be much more heterogeneous between batches. CAM ComBat will be implemented in the following way.

#### Step 1

Standardize the data similarly to ComBat where residuals are achieved after estimating and removing the biological variabilities and rescaled. Letting *Z*_*ijv*_ represent the standardized data,

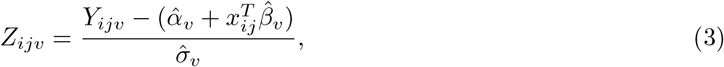

where 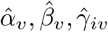 are the feature-wise ordinary least squares estimates and 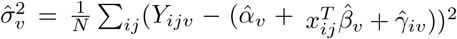 with *N=Σ*_*i*_*n*_*i*_

#### Step 2

Let *Z*_*ij*_ = (*Z*_*ij*1_, …, *Z*_*ijV*_)^*T*^ and assume 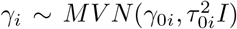. Using the formula for the expectation of the posterior distribution of *γ*_*i*_, calculate the posterior mean for *γ*_*i*_ (derivations shown in Appendix):

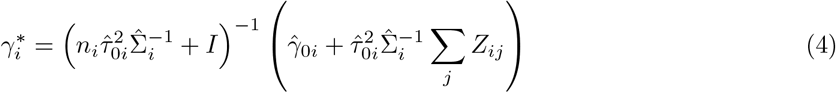

in which 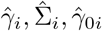, and 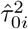 are their corresponding method of moments estimates.

#### Step 3

Assume that 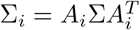. Using the eigendecomposition of 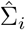and 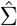, the sample covariance estimate of Σ, *A*_*i*_ can be estimated as follows (see calculations in Appendix):

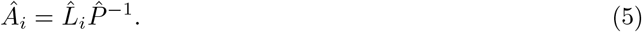

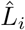 and 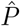are calculated as

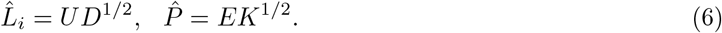

where *U* is the matrix of eigenvectors of 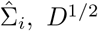 is the diagonal matrix of the square root of the eigenvalues of 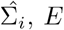 is the matrix of eigenvectors of 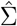, and 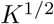 is the diagonal matrix of the square root of the eigenvalues of 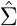.

#### Step 4

Adjust the data as follows to get the final data *M*_*ij*_:

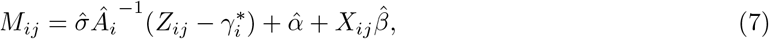

where 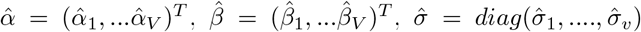, and *X*_*ij*_ is the matrix of covariate information.

### 2.2 SIB ComBat for High Dimensional Vertex Level Data

In practice, applying CAM ComBat to a high dimension of features can be computationally intensive due to the estimation of the overall and within-site covariance matrices across all features of interest. If we are interested in harmonizing vertex-level neuroimaging data, which can have hundreds of thousands of features, CAM ComBat becomes impractical. To address this limitation, we developed an extension of the method, Spatially-informed Iterative Block (SIB) ComBat, that can alternatively be used.

SIB ComBat implements an iterative process for harmonization within pre-defined clusters of features. Clusters of features are harmonized separately using CAM ComBat in a given iteration and the process iterates through different partition of features. In each iteration, feature clusters are harmonized separately using CAM ComBat, and the procedure cycles through multiple feature partitions across iterations. Optimal cluster assignments are those in which features exhibit strong within-cluster correlations while remaining relatively independent across clusters. Examples of such cluster assignments are shown in Figure 1, which visualizes clusters determined using multi-scale structural imaging covariance (MuSIC) atlases that will be discussed later [Wen et al., 2023]. Applying CAM ComBat on subsets of features instead of all features helps to make this method more computationally feasible. The iterative harmonization process enables us to account for correlation between more pairs of features by harmonizing different groups of features in each iteration, which allows us to better adjust for overall covariance site effects in the data. The steps of the algorithm can be found in algorithm 1.

**Figure 1:**
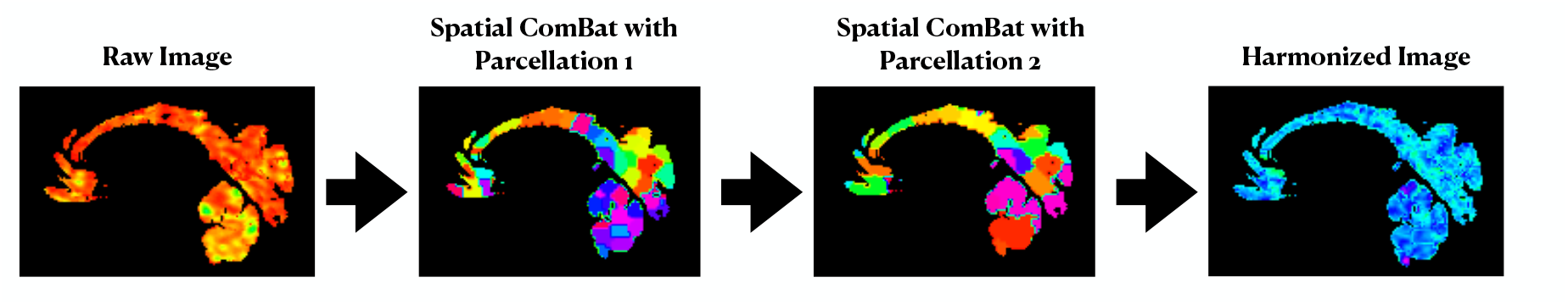
Diagram visualizing the SIB ComBat harmonization process.

#### Algorithm 1

SIB ComBat

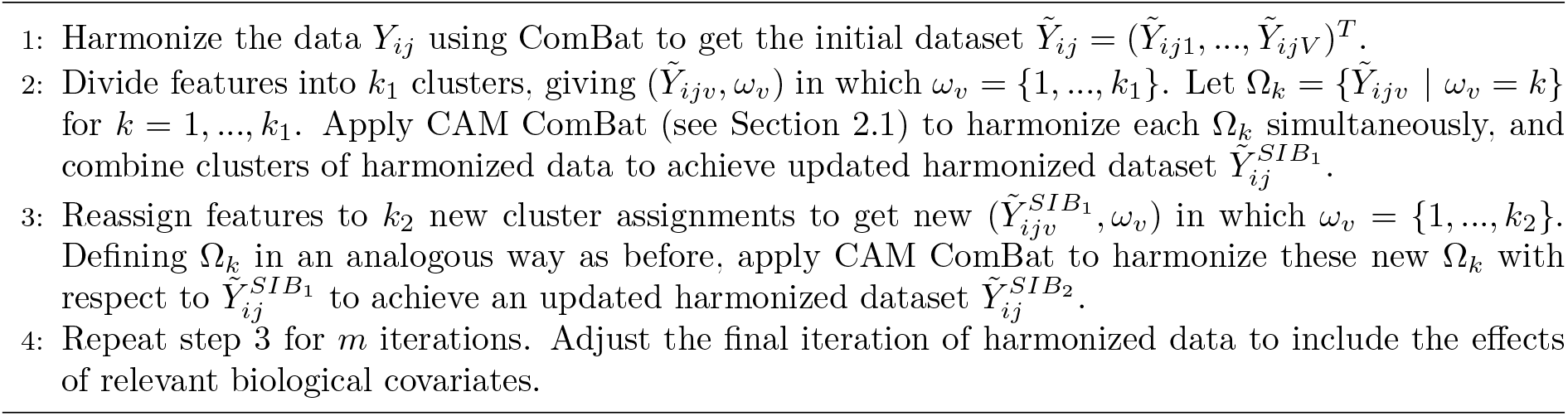

## 3 Simulations

### 3.1 Experiment Set Up

We first conducted simulation experiments to compare the performance of CAM ComBat and SIB ComBat to ComBat and CovBat. To do so, data were simulated from *n* individuals with 100 features each based on the following model, in which *{e*_*ij*1_, *…, e*_*ijv*_*}*^*T*^ ∼ *MV N* (**0**, Σ_*i*_):

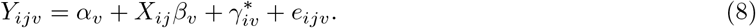

Individuals were generated to have a binary covariate and belong to one of two batches, with each batch having an equal number of individuals. Binary covariates were randomly generated for each individual from a Bernoulli(0.5) distribution. *α*_*v*_ and *β*_*v*_ were generated from a *N* (0, 1) distribution, and 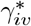for each batch was generated from a *N* (4, 1) distribution. Σ_*i*_ for each batch was generated from an InverseWishart distribution with scale matrix Ψ and degrees of freedom *v*.

Datasets were simulated under various scenarios to evaluate the performance of the harmonization methods in different settings. Comparisons were made between total sample sizes of 100 and 200, and with sets of batch-specific covariance matrices with varying magnitudes of difference, as calculated using the Frobenius distance. To achieve different levels of site effects, each pair of Σ_*i*_ matrices were generated from an inverse Wishart distribution with the same scale matrix Ψ but different degrees of freedom *v*, using the *riwishart* function from the *MCMCpack* package in R [Martin et al., 2011]. Larger degrees of freedom were used to generate Σ_*i*_ that were less variable, resulting in more similar covariance matrices. Batch-specific covariance matrices were specifically generated with *v* values of 108, 106, and 104.5. The resulting Frobenius distances of the respective pairs of covariance matrices were 617.04, 1641.62, and 3532.028, which will represent small, medium, and large matrix differences.

The simulated data were harmonized using ComBat, CovBat, CAM ComBat, and SIB ComBat. For each simulated dataset, SIB ComBat was fixed to apply two iterations of harmonization for all datasets. Each iteration harmonized two randomly determined clusters of features, with 50 features in each cluster. To compare the performance of each method, data from 50 individuals were randomly chosen as the training set, and data from another 50 subjects were randomly selected as the validation set. A random forest algorithm was trained to detect batch using the training data with the *randomForest* function from the *randomForest* package in R [Liaw and Wiener, 2002]. To evaluate the predictive performance of the random forest model on the validation data for batch detection, we calculated the area under the curve (AUC), using the *auc* function from the *pROC* package in R [Robin et al., 2011]. This was repeated over ten random permutations of training and validation sets, and the resulting AUC’s were averaged. The same analysis was conducted for classifying the binary covariate level. The overall setup was repeated for 100 simulated datasets, and results were visualized with side-by-side boxplots.

### 3.2 Simulation Results

With a smaller difference in the covariances between batches, the total sample size affected the relative performance of every harmonization method in mitigating batch effects. When the total sample size was 100, all methods had relatively comparable AUC’s for batch detection, with the lowest AUC achieved using CovBat and the highest AUC achieved using SIB ComBat on average. In contrast, when the sample size was 200, Spatial and SIB ComBat had notably lower AUC values for detecting batch on average and showed more notable improvements in mitigating batch effects compared to the other methods. This shows that Spatial and SIB ComBat better mitigate batch detection with a larger sample size. Given the comparable AUC’s between Spatial and SIB ComBat when the total sample size was 200, this indicates that SIB ComBat may be a good alternative to CAM ComBat if the sample size is sufficient. However, regardless of the sample size, all methods were relatively sufficient to mitigate batch effects, which makes sense in this setup since the covariances between batches are more similar, and thus, the batch effects are more directly tied to the means and variances (see Figure 2).

**Figure 2:**
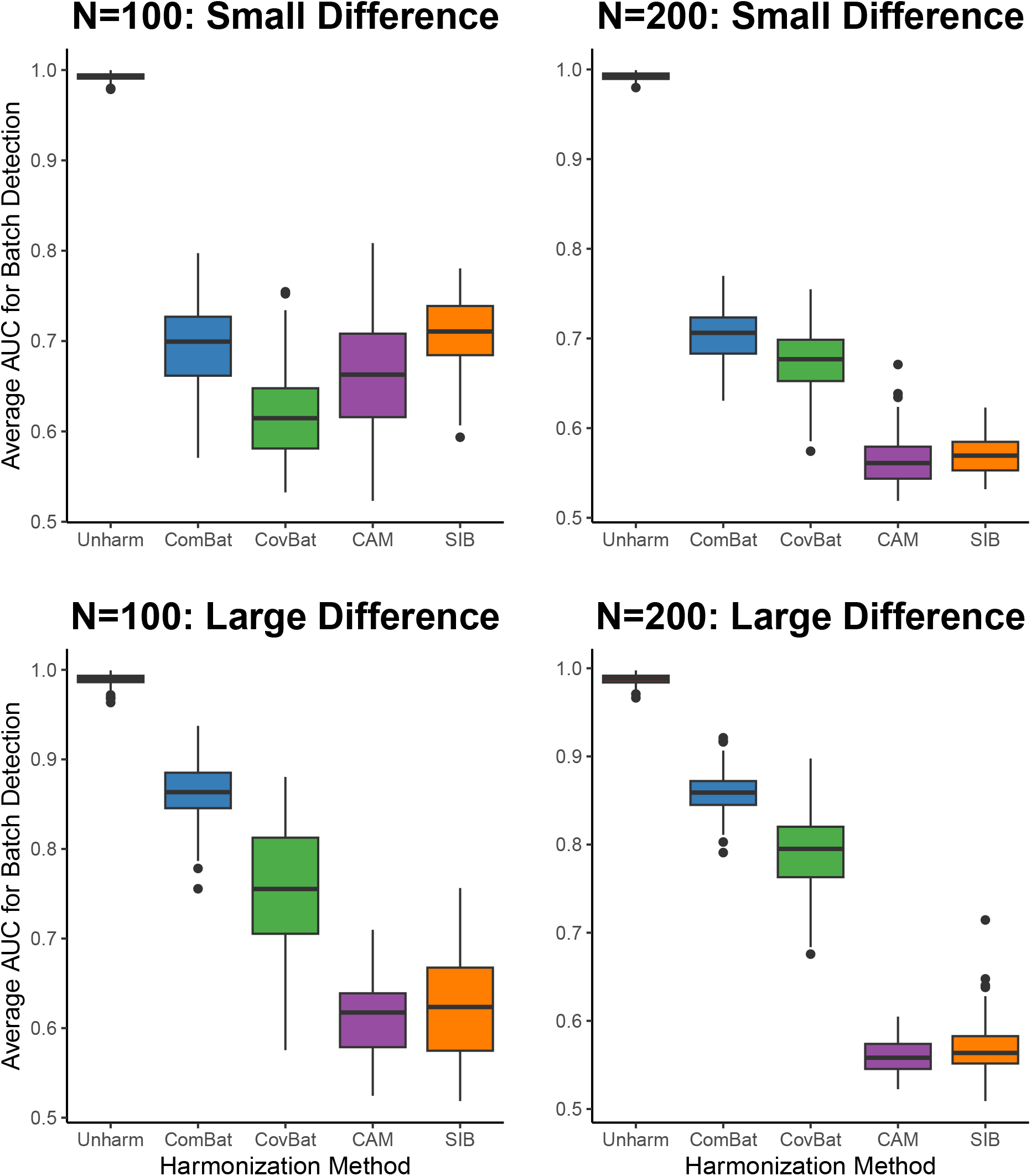
Side-by-side boxplots comparing AUC for detecting batch between ComBat, CovBat, CAM ComBat, and SIB ComBat. Comparisons done between N=100 (left) and N=200 (right) and between smaller (top row) versus larger (bottom row) differences between covariances.

If the difference in Σ_*i*_ between batches was large, CAM ComBat was able to consistently remove batch effects, regardless of the total sample size. It decreased the mean AUC for detecting batch to a value below 0.7, which was not seen for ComBat and CovBat. SIB ComBat also demonstrated a comparable performance in mitigating the detection of batch, achieving AUC’s that were comparable to those of CAM ComBat. In general, these results support SIB ComBat as a potential alternative to CAM ComBat when the dimension of features is high. For both Spatial and SIB ComBat, there was a particularly large improvement when the total sample size was 200, further emphasizing the benefit of larger sample sizes found previously. These results suggest that when there is more heterogeneity of covariances between batches, Spatial and SIB ComBat will perform the best in removing batch effects, especially when the sample size is larger (see Figure 2).

In addition, under both magnitudes of matrix differences, every harmonization method was able to preserve the ability of classifying the binary covariate with similar or higher AUC’s compared to the unharmonized data. This indicates that every harmonization method was able to maintain at least as strong a relationship with relevant biological covariates as found in the original data. We do notice a particularly large increase in the AUC’s for binary covariate classification when using SIB ComBat compared to the other methods, regardless of the total sample size. This suggests that SIB ComBat can magnify the strength of a relationship with respect to a biological covariate (see Figure 3).

**Figure 3:**
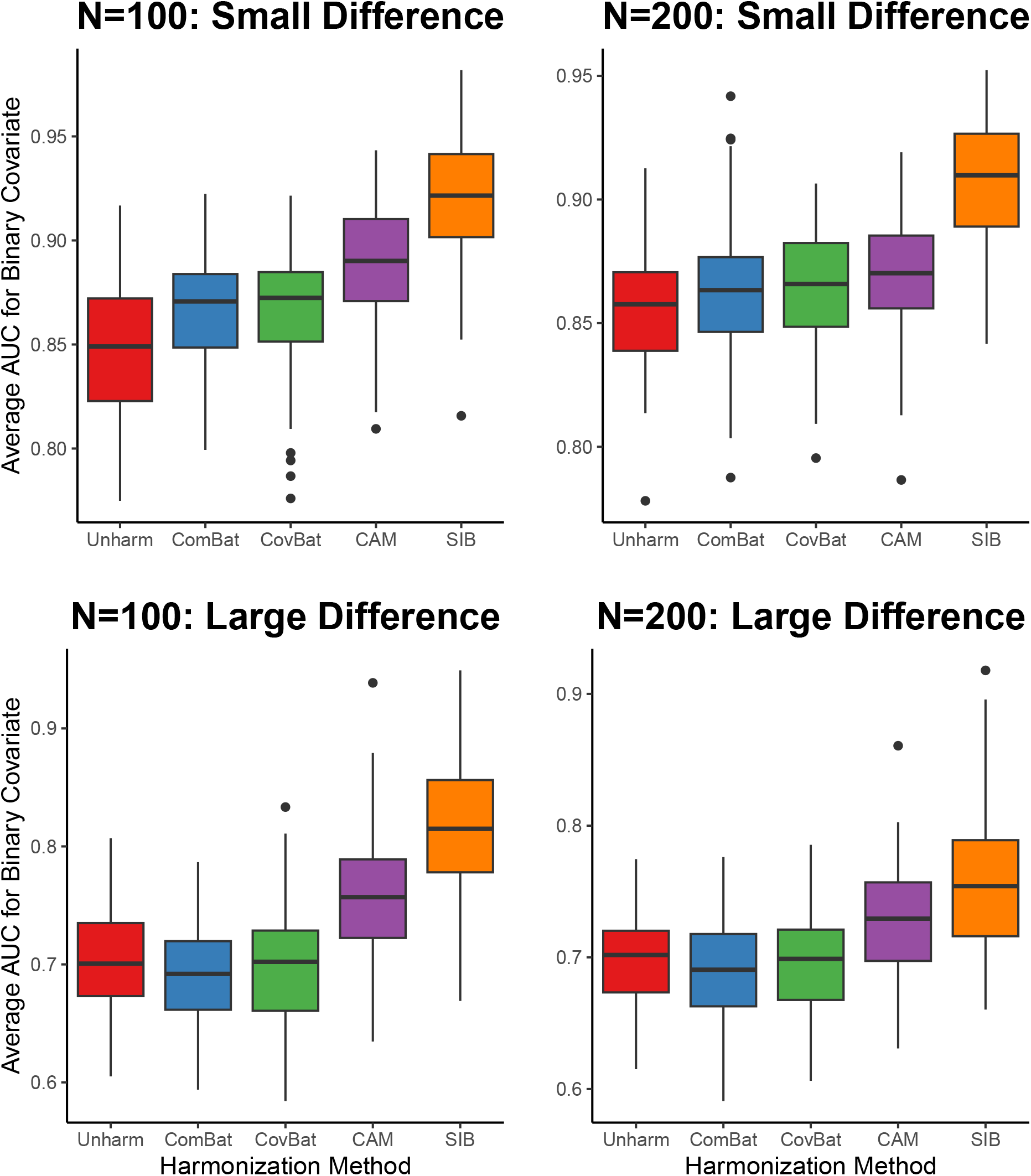
Side-by-side boxplots comparing AUC for detecting binary covariate level between ComBat, CovBat, CAM ComBat, and SIB ComBat. Comparisons done between N=100 (left) and N=200 (right) and between smaller (top row) versus larger (bottom row) differences between covariances.

## 4 Real Data Application

### 4.1 Data

We applied the proposed methods to voxel-wise RAVENS gray matter density maps from two studies of the iSTAGING consortium: the Harvard Aging Brain Study (HABS) and the Open Access Series of Imaging Studies (OASIS). Scans were obtained from HABS using the Siemens Magnetom Tim Trio 3T scanner at Massachussett’s General Hospital, while those from OASIS were obtained with the Siemens MPRAGE sequence at 1.5 or 3 T [Dagley et al., 2017, LaMontagne et al., 2019]. T1-weighted MRI scans were processed using an in-house automated pipeline. Each T1 scan was first reoriented, correcting for intensity inhomogeneities. Multi-atlas based skull-stripping using a large set of reference atlases was used for the extraction of brain from extra cranial tissues. Skull-stripped T1 images were segmented into a set of anatomical regions of interest (ROIs) (145 single ROIs – 119 Gray Matter (GM), 20 White Matter (WM) and 6 ventricular regions) using a multi-atlas label fusion method, MUSE. This is an atlas guided segmentation of anatomical regions that involves warping an atlas into the subject space to establish spatial correspondence between two images and then transferring atlas ROI labels to the subject space using two deformable registration algorithms, DRAMMS and ANT [Doshi et al., 2016, Ou et al., 2011, Avants et al., 2008]. We used label fusion wherein labels from multiple warped atlases were fused together to determine final labels. RAVENS tissue density maps were generated after segmenting the brain into GM, WM, and Cerebrospinal Fluid (CSF) followed by high dimensional warping to a common template [Davatzikos et al., 2001]. For our analysis, we focused on a subset of 500 individuals, half of which came from HABS, and half of which came from OASIS.

Our final processed dataset consists of 641,838 features measuring RAVENS gray matter density maps derived from structural MRI, where the intensity value at each voxel represents the amount of gray matter present in that particular region of the brain [Wen et al., 2023]. On average, subjects were 71.6 years old, with 61.6% of individuals reporting being female and 85.2% of individuals reporting being Caucasian. Additionally, 1.6% of subjects were reported to have some form of cognitive impairment, including mild cognitive impairment, Alzheimer’s disease, and dementia. More specific overall and study-specific demographic information are provided in Table 1. As illustrated in Figure 6, the average map of the voxel-wise intensity measurements from the two studies demonstrate visual differences in the mean measurements of voxels depending if the subject was from the HABS or OASIS studies. We are specifically interested in harmonizing such data to eliminate study-specific effects.

**Table 1:** Table comparing age, sex, deep learning intracranial volume (DLICV), race (% Caucasian), and % with reported cognitive impairment (CI) such as dementia and Alzheimer’s disease (DX) between the subset of HABS and OASIS subjects. For age and DLICV, we conducted two-sample t-tests and reported the means and standard deviations. Fisher’s exact tests were performed for sex, DX, and race, with the respective percentages reported.

|  | Overall | OASIS | HABS | P-Value |
| --- | --- | --- | --- | --- |
| Age | 71.6 (8.7) | 69.4 (10.2) | 73.8 (6.2) | 1.31e-08 |
| DLICV ($\times 10^6$) | 1.39 (0.144) | 1.39 (0.147) | 1.40 (0.140) | 0.38 |
| Sex (% Female) | 61.6 | 60.8 | 62.4 | 0.78 |
| DX (% CI) | 1.6 | 2.0 | 1.2 | 0.72 |
| Race (% Caucasian) | 85.2 | 89.2 | 81.2 | 0.02 |
| N | 500 | 250 | 250 |  |

### 4.2 Analysis

The data were harmonized using ComBat, CovBat, and SIB ComBat. In the harmonization process, we account for age, sex, intracranial volume at baseline, cognitive impairment, and race. When using SIB ComBat, two iterations of harmonization were conducted, where clusters for each iteration were determined using a MuSIC atlas. These atlases were derived from data using stochastic orthogonally projective non-negative matrix factorization to capture patterns in structural covariance across brain regions at various scales. Non-negative matrix factorization decomposes complex neuroimaging data into a sparse, part-based brain representation by projecting it onto a few components, and has been lauded for its improved clinical interpretability compared to other unsupervised methods [Wen et al., 2023]. For the first iteration of SIB ComBat harmonization, features were clustered based on the 512 MuSIC atlas, while features were clustered based on the 256 MuSIC atlas for the second iteration of harmonization.

To evaluate the performance of these harmonization methods and avoid overfitting due to the large dimensions of features, we first calculated the average feature measurement within a cluster across 256 clusters, determined using the 256 MuSIC atlas. Data from 100 individuals were randomly chosen as the training set, and data from another 100 subjects were randomly selected as the validation set. Random forest algorithms were trained to detect study site and classify sex using the training data. To evaluate the predictive performance of the random forest models in the validation data for study site detection and sex classification, we calculated the AUC [Robin et al., 2011]. This was repeated over 100 random permutations of training and validation sets, and results were visualized with side-by-side boxplots.

In addition to this analysis, we conducted a voxel-based morphometry (VBM) analysis, in which we fit a linear model for gray matter density maps at each voxel, including age, sex, intracranial volume at baseline, cognitive impairment, and race as covariates. We extracted the t-statistics testing for the significance of age and mapped those statistics back to images of the brain. This analysis was conducted for the unharmonized data and the SIB-ComBat harmonized data to analyze the effects of harmonization on the associations with age. Furthermore, we calculated the respective t-statistics if we only used data from the HABS or OASIS studies and analyzed their concordance using a concordance at the top (CAT) plot across different harmonization methods [Irizarry et al., 2005]. We also conducted another analysis similar to before in which we fit a linear model for gray matter density maps at each voxel, and included study site along with age, sex, intracranial volume at baseline, cognitive impairment, and race as covariates. The t-statistic for study site was computed across all voxels and the distributions of these statistics were compared between different harmonized datasets.

### 4.3 Results

Across all methods, SIB ComBat was the most effective in removing study-specific effects, being the only method to decrease the mean AUC for detecting study site to a value below 0.7. SIB ComBat was also able to magnify the association with respect to sex, as seen by the higher AUC’s for classifying sex compared to the other harmonization methods (see Figure 4). Furthermore, looking at the maps of the VBM t-statistics before and after SIB ComBat was applied, they appear pretty similar to each other.

**Figure 4:**
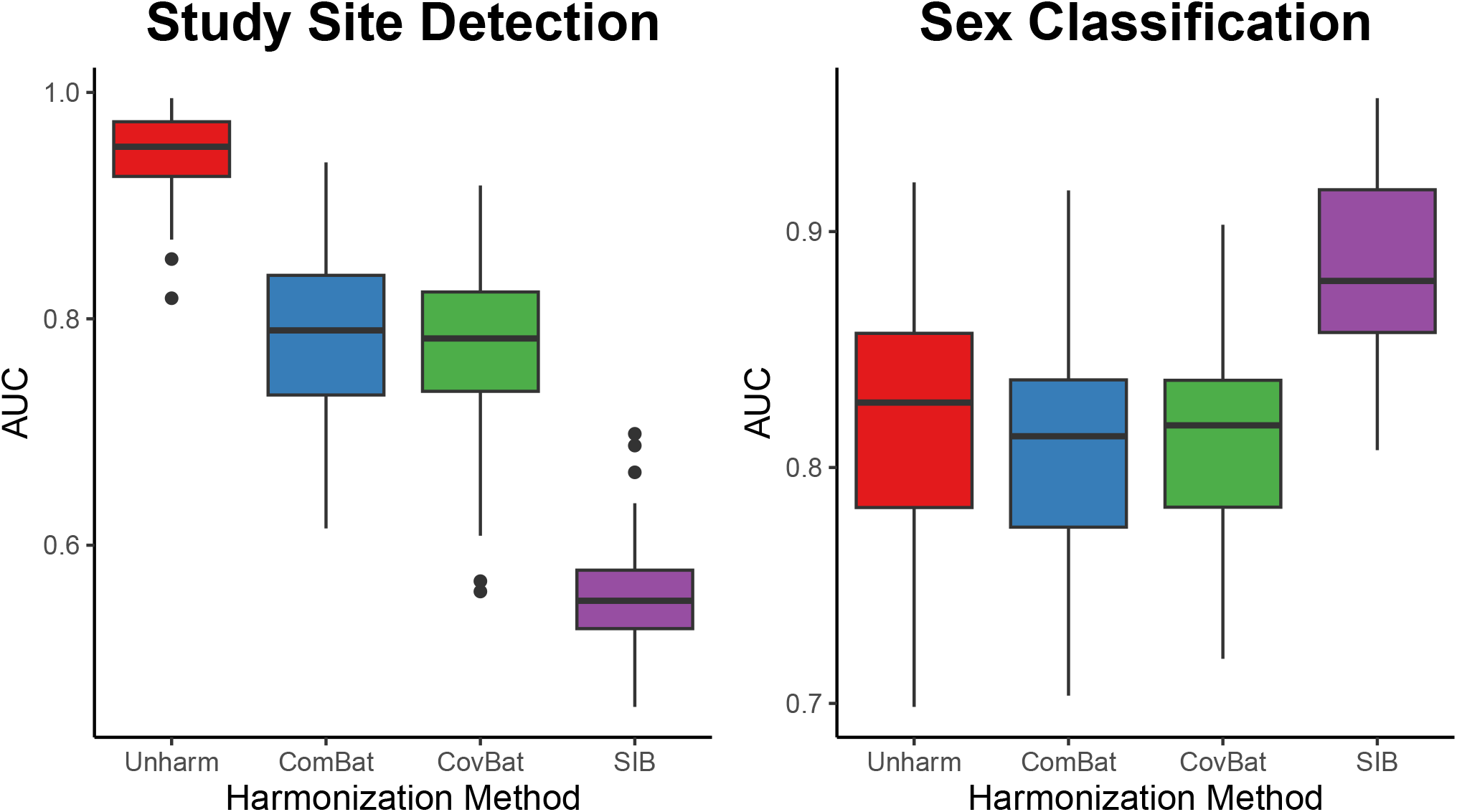
Side-by-side boxplots comparing AUC for detecting study site (HABS or OASIS) and classifying sex between ComBat, CovBat, and SIB ComBat using multi-site voxel-level intensity measurements.

There are no visually significant differences in both the relative magnitudes of the test statistics and their spatial location within the brain, indicating that the association with age was generally preserved after harmonization (see Figure 5). Maps of the average voxel intensities between HABS and OASIS participants showed less noticeable differences after harmonizing with SIB ComBat compared to the original data (see Figure 6). This finding was further supported by the low t-statistics for study site, which were generally comparable across harmonization methods (Figure 7). There was also uniformly better concordance between the HABS and OASIS t-statistics testing for the association with age after SIB ComBat was applied, and had the best concordance across different harmonization methods (see Figure 8). This indicates that SIB ComBat harmonization was able to achieve better agreement between sites in voxels significantly associated with age compared to other unharmonized and harmonized datasets.

**Figure 5:**
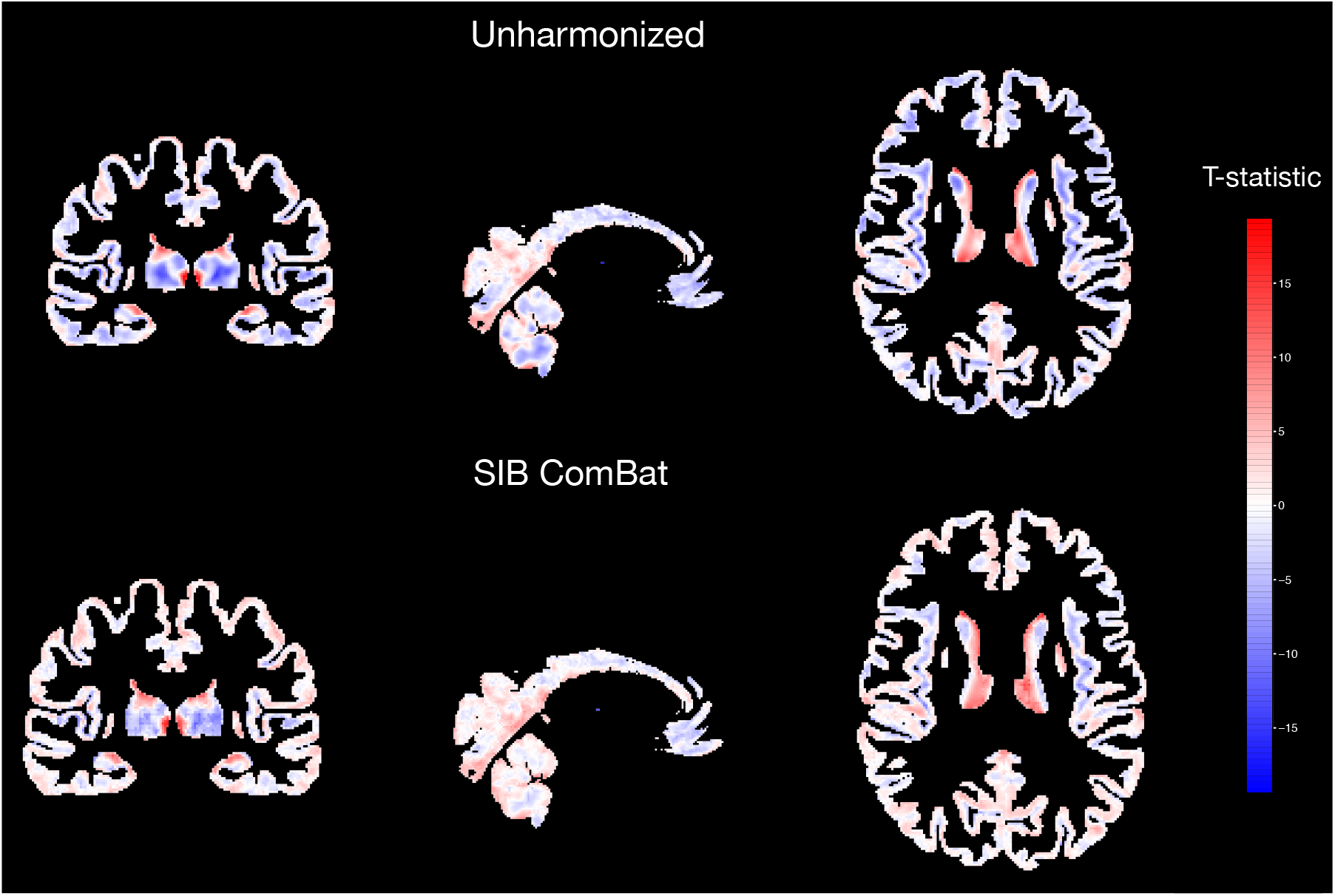
Voxel-based morphometry maps illustrating magnitude of t-statistics when testing for significance of age, conditioned on sex, deep learning intracranial volume, race, and cognitive impairment, using unharmonized and SIB-ComBat harmonized data.

**Figure 6:**
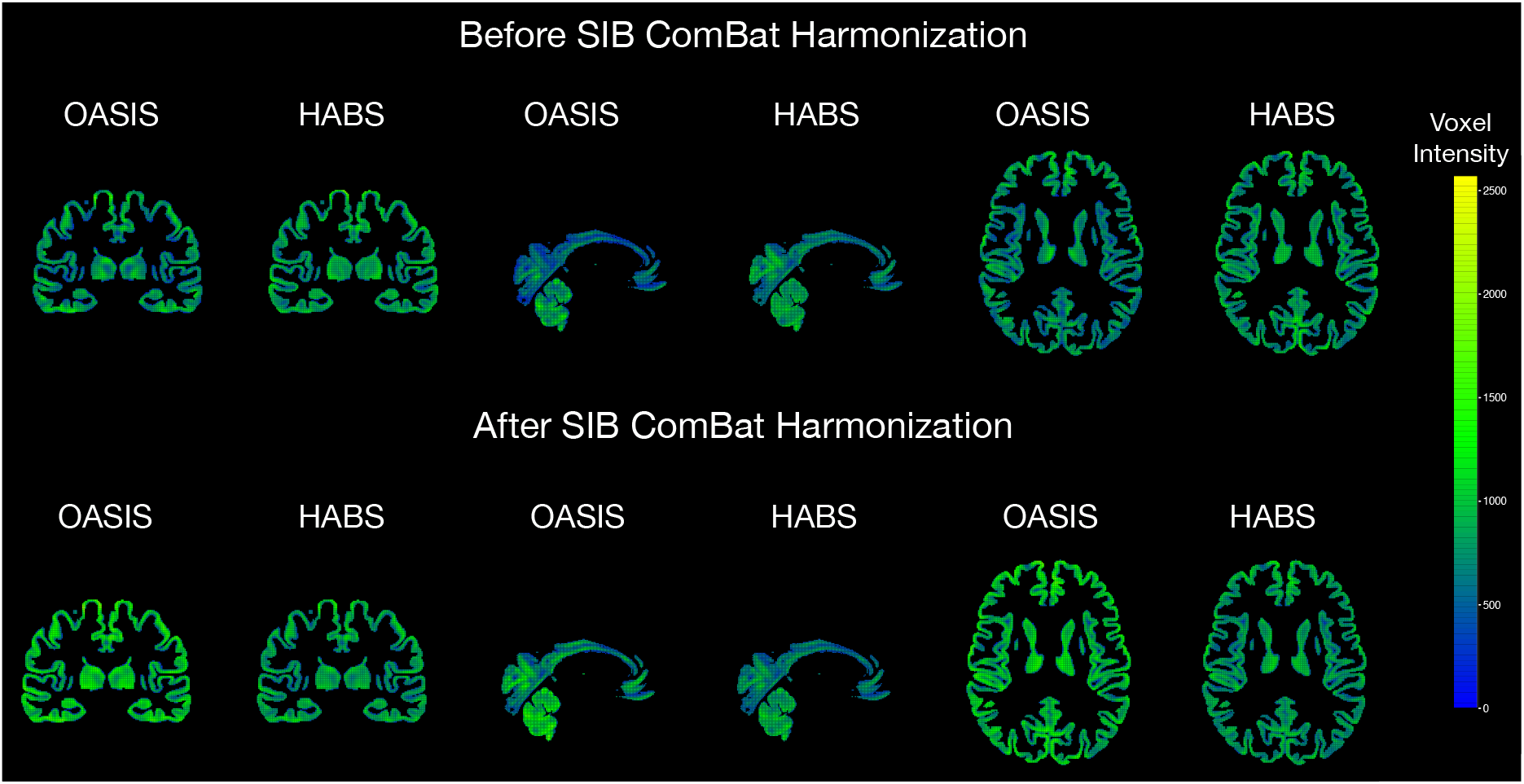
Maps of the brain representing average voxel-level intensity measurements if individuals were from OASIS or HABS before and after harmonizing with SIB ComBat.

**Figure 7:**
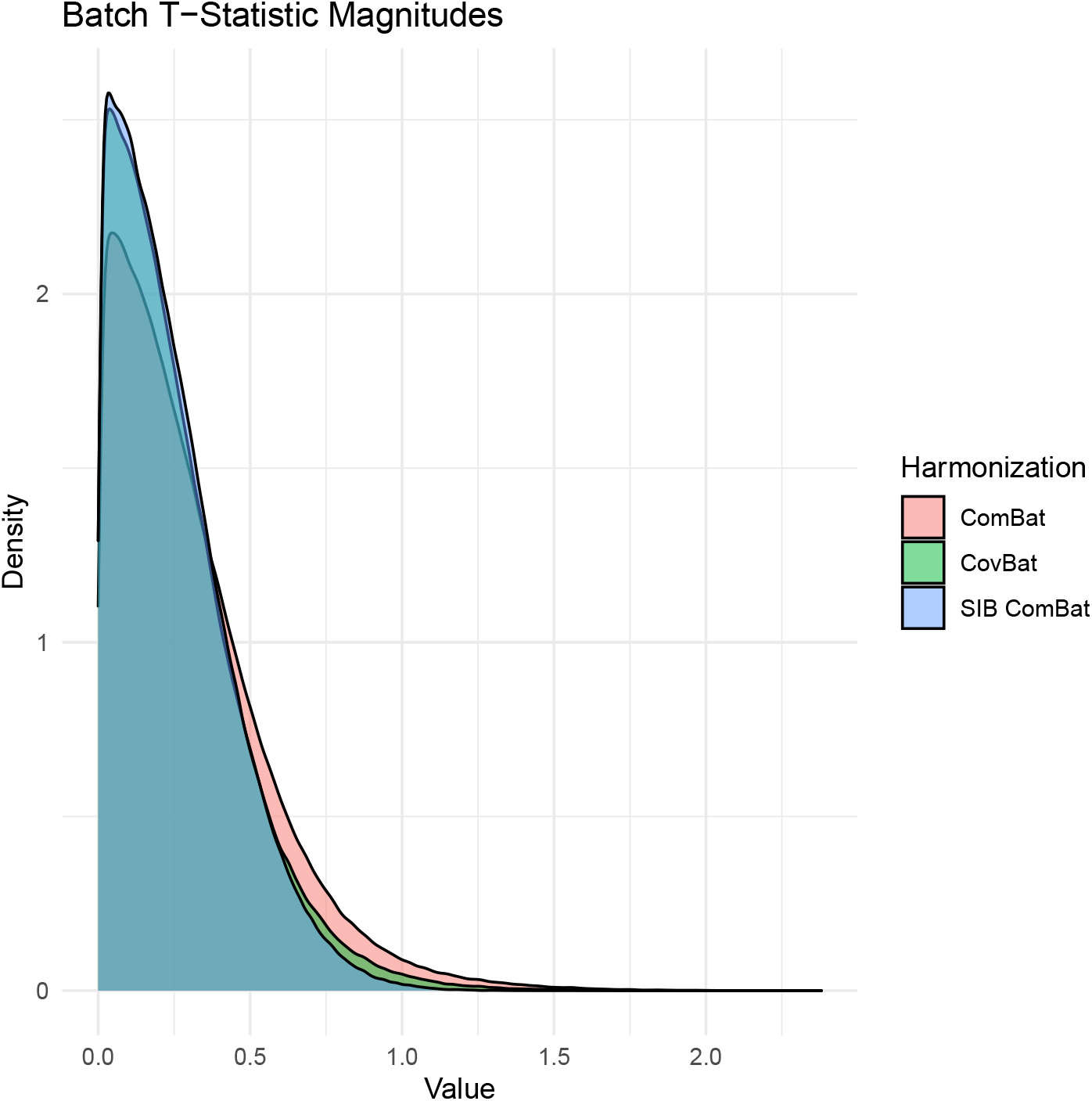
Densities of t-statistic magnitudes for study site for different harmonized data sets, after adjusting for age, sex, deep learning intracranial volume, race, and cognitive impairment.

**Figure 8:**
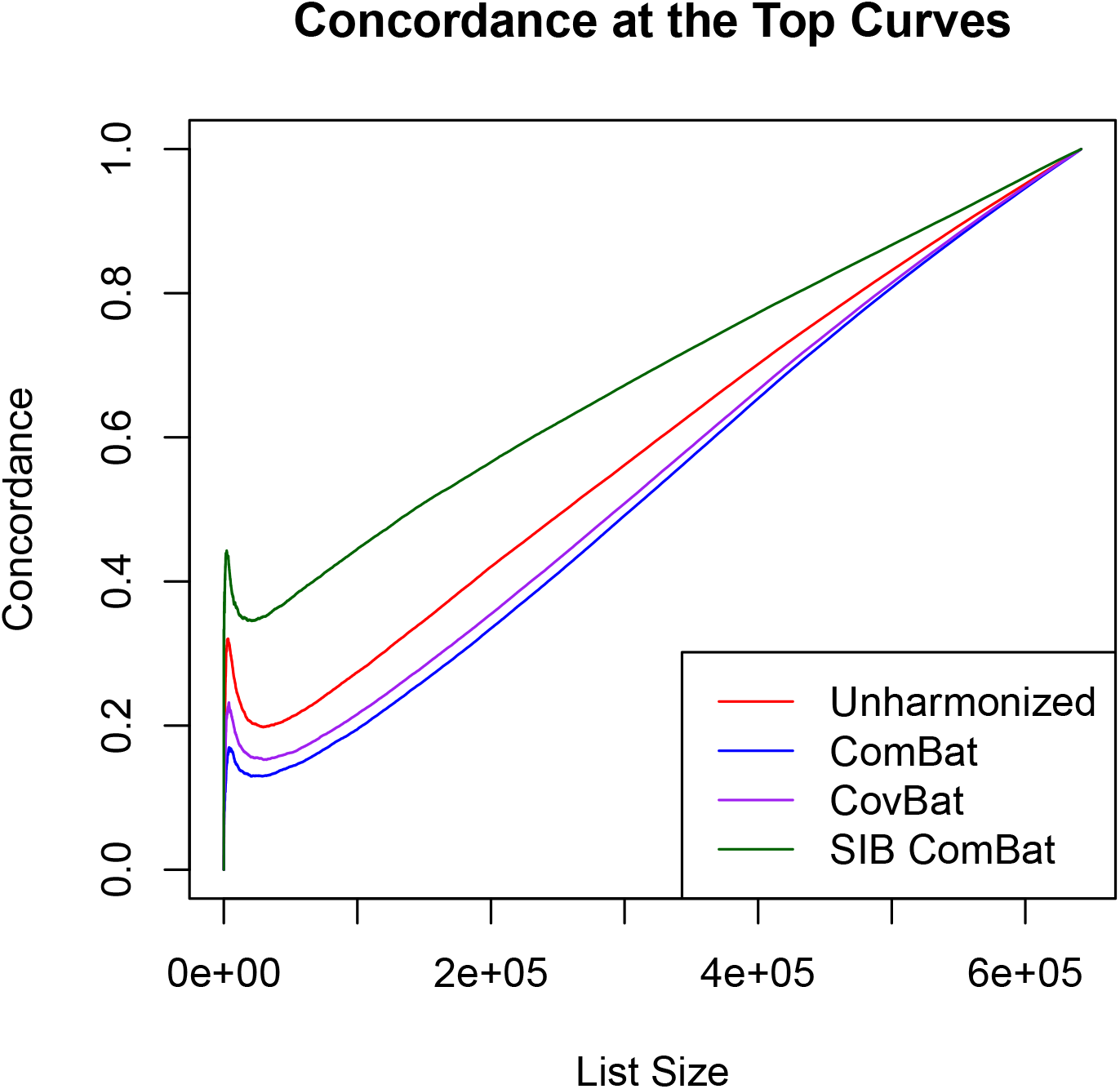
Concordance at the top (CAT) plot comparing the concordance of t-statistics for age between HABS and OASIS studies with unharmonized, ComBat-harmonized, CovBat-harmonized, and SIB ComBat-harmonized datasets.

**Figure 9:**
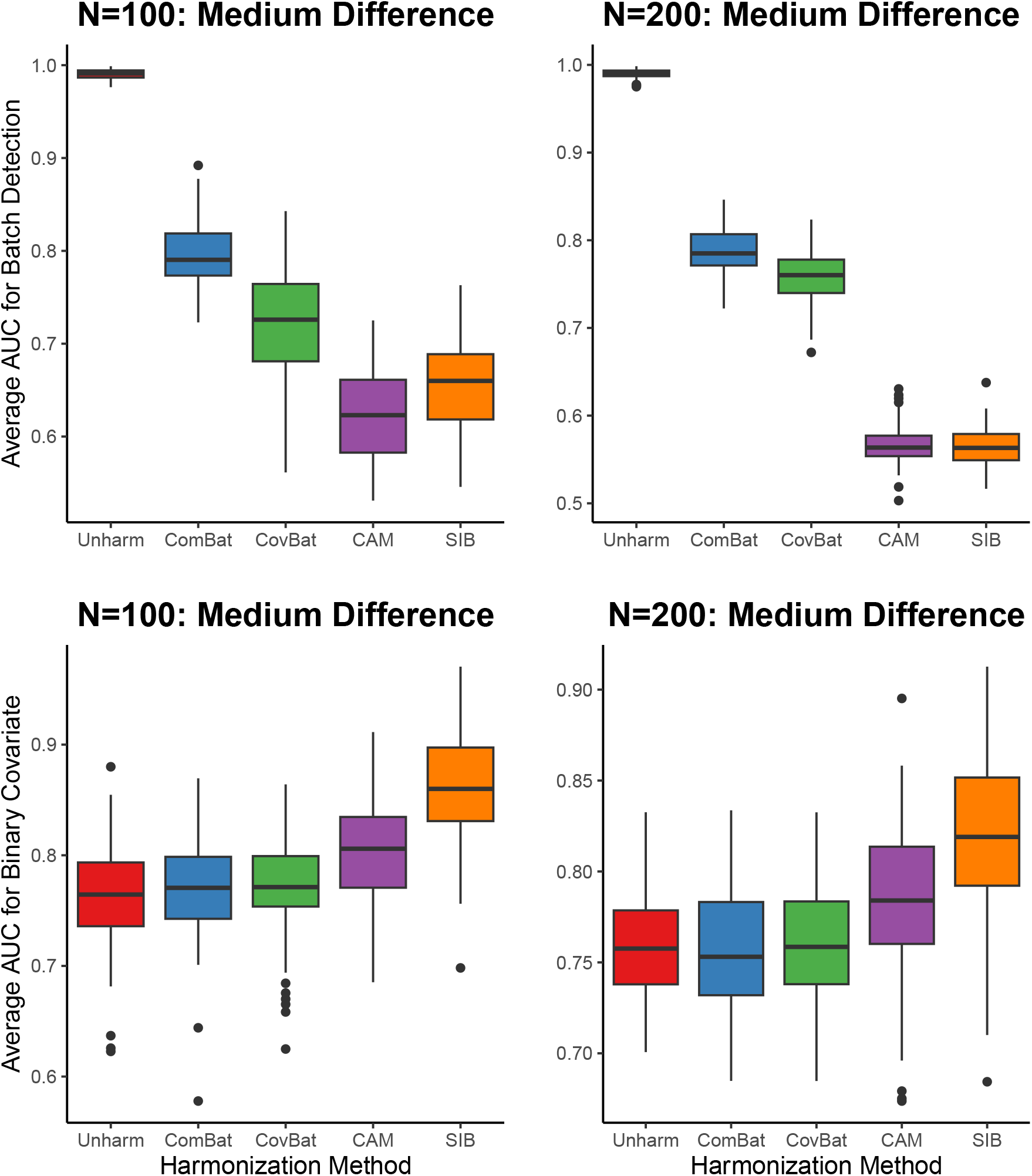
Side-by-side boxplots comparing AUC for detecting batch (top row) and binary covariate level (bottom row) between ComBat, CovBat, CAM ComBat, and SIB ComBat. Comparisons done between N=100 (left) and N=200 (right) for medium difference between covariances.

Interestingly, the concordances for the datasets harmonized with ComBat and CovBat were uniformly lower compared to the unharmonized data. When looking into this result, we noticed that some features showed differing distributions of intensities between HABS and OASIS individuals. Specifically, we saw a larger proportion of individuals from OASIS having relatively lower intensity values. ComBat and CovBat further emphasized these distributional differences, resulting in t-statistics for age that differed more between sites and subsequently worse concordance. This behavior makes sense given ComBat and CovBat both remove site-specific variability by rescaling the residuals with respect to the site-specific variance. While CovBat also shifts and rescales the principal component (PC) scores corresponding to the PC decomposition of the full covariance matrix, the difference in the distribution of intensities between HABS and OASIS individuals is still notable. In CAM (and subsequently SIB) ComBat, the individual’s vector of residuals is multiplied by the matrix 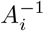, which is estimated using the eigendecompositions of the full and site-specific covariance matrices. Doing so enables us to leverage more information across features, resulting in a stronger adjustment to the residuals that minimizes this distributional difference and gives stronger concordance. An example of this comparison can be found in Figure 10 in the Appendix. All in all, SIB ComBat had the best overall performance in significantly reducing the detection of site while generally maintaining the associations found with respect to age and sex in the original data.

**Figure 10:**
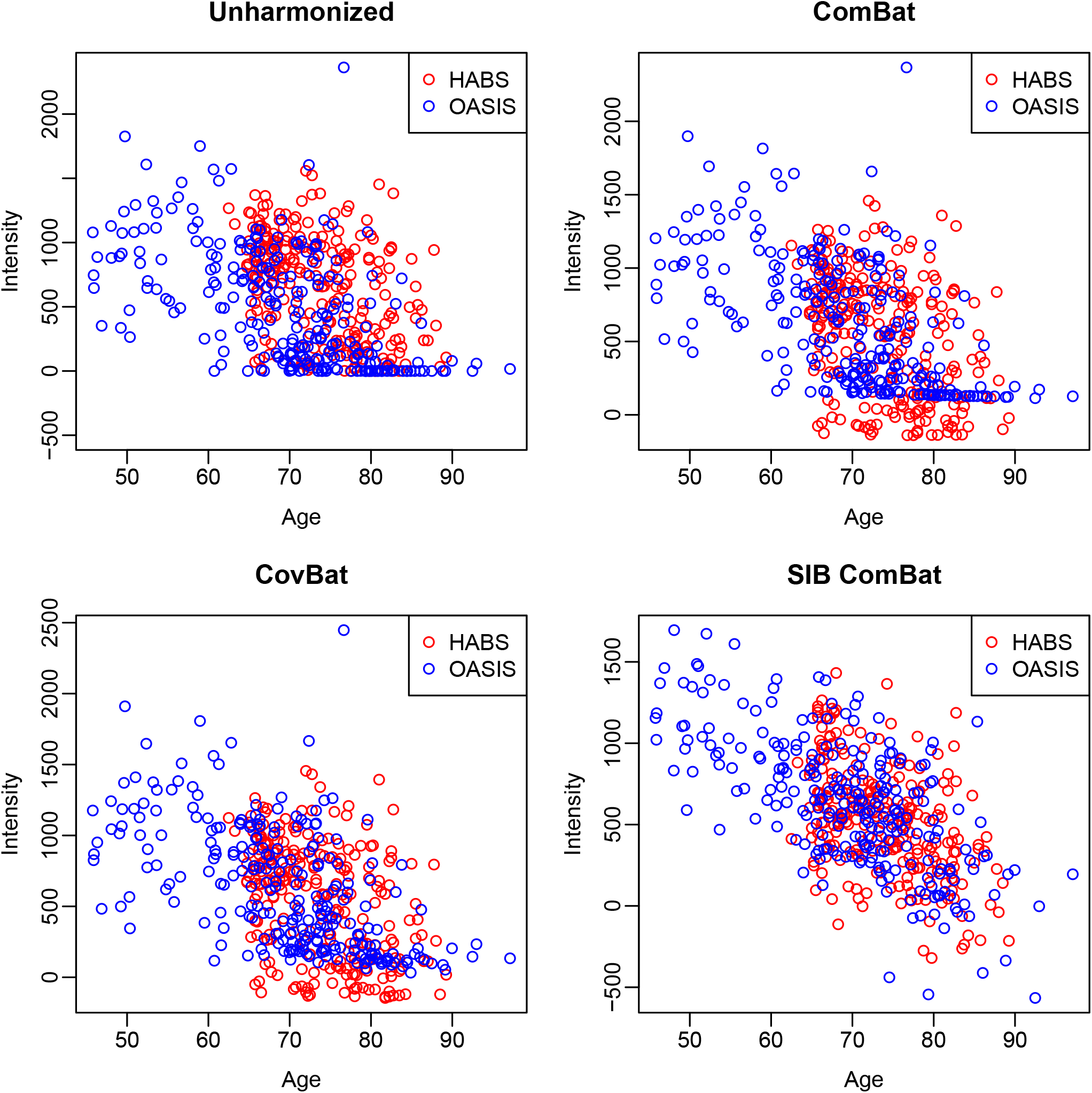
Scatter plots visualizing the association between the intensity of a given voxel-level feature and age across different harmonized datasets. Associations are compared between individuals from HABS against those from OASIS.

**Figure 11:**
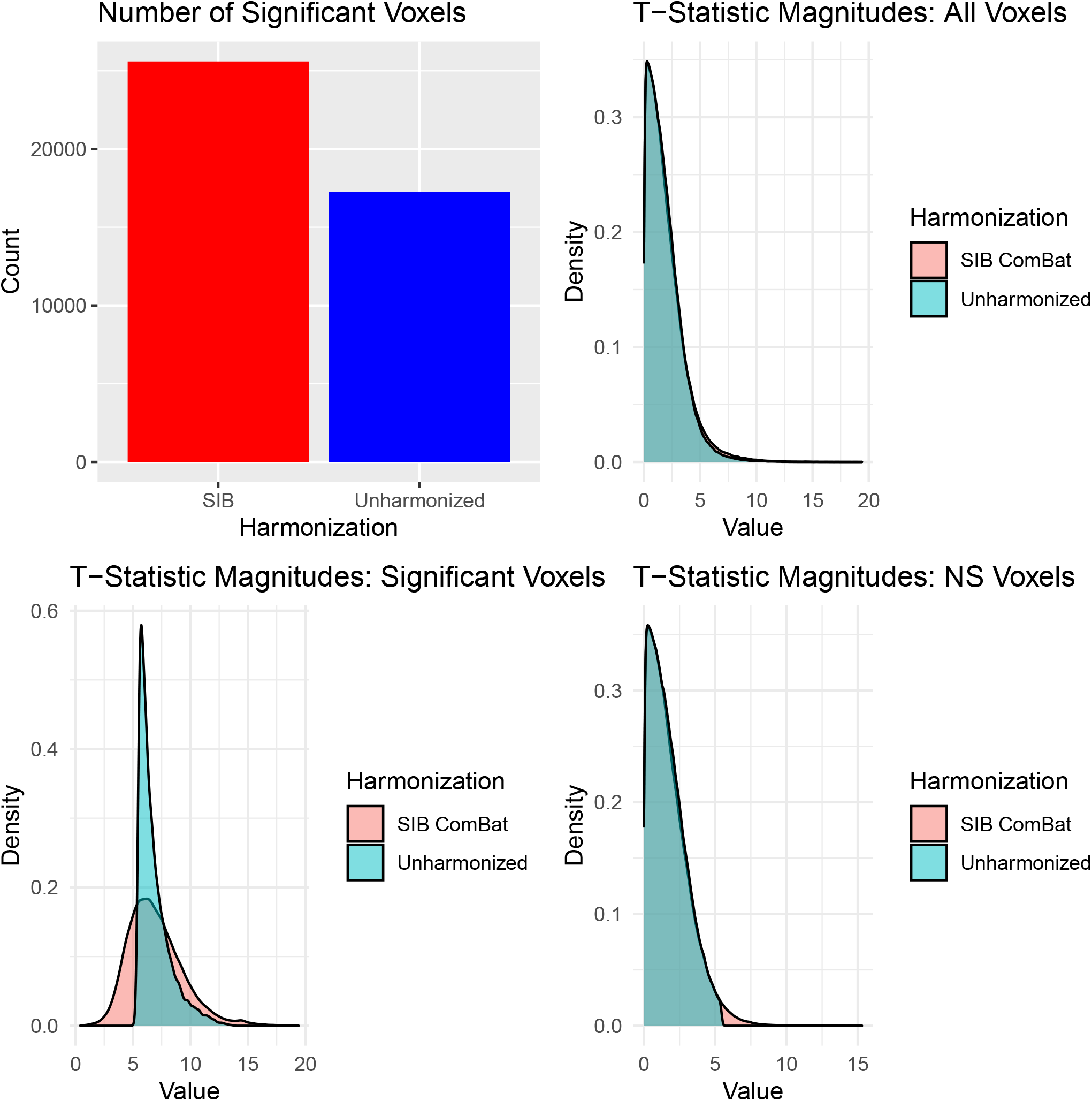
Plots showing the number of significant voxels for age (upper left) and the distributions of age t-statistic magnitudes across all voxels (upper right) / originally significant voxels (lower left) / originally not significant (NS) voxels (lower right) between SIB ComBat-harmonized and unharmonized data.

**Figure 12:**
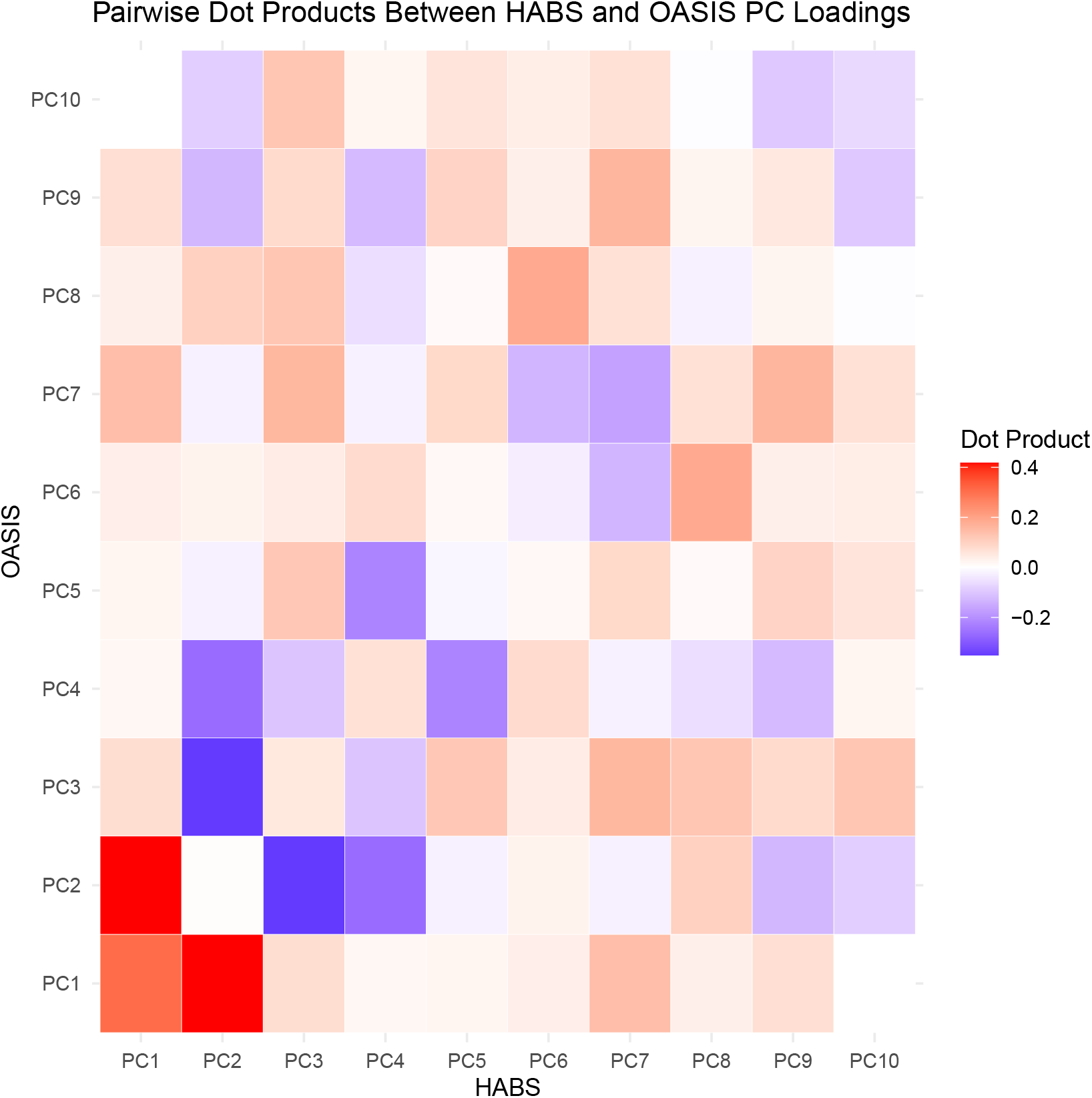
Heatmap that illustrates the pairwise similarity of the first ten principal component eigenvectors between two study sites, HABS and OASIS, by calculating their respective dot product. In this figure, most of the dot products between eigenvectors were close to 0, demonstrating strong dissimilarity.

## 5 Discussion

Through various simulation experiments, compared to ComBat and CovBat, CAM ComBat was the most robust in removing batch effects under varying levels of heterogeneity between the batch-specific covariance matrices, with a notably better performance when the heterogeneity between batches was large. It demonstrated superior performance with a larger sample size, which implies that sample size affects how well CAM ComBat can harmonize the data. CAM ComBat also maintained associations with the simulated binary covariate to a similar degree as ComBat and CovBat. However, it may not be a practical method to implement when harmonizing a large number of features. We thus developed SIB ComBat, which expands upon CAM ComBat to accommodate these higher dimensional scenarios. In our simulation experiments, SIB ComBat proved to be equally effective in mitigating batch effects while magnifying the strength of the binary covariate association. When comparing the performance of SIB ComBat to ComBat and CovBat for harmonizing multi-site voxel intensity measurements from the iSTAGING consortium, SIB ComBat was superior in removing site-specific effects while maintaining the strength of the association found with respect to sex and age in the original data.

Although we developed SIB ComBat to make it more accommodating in higher dimensional situations, there are still practical limitations in its implementation. Despite the improvements relative to CAM ComBat, SIB ComBat can still take a long time to run and use a lot of memory. This method benefits from parallelizing the cluster-specific harmonization. Although this is reasonable if using a cluster with multiple nodes, it may be less feasible on a local laptop. With our implementation of SIB ComBat using a total of eight cores, it took over 36 hours to run and used over 120 GB of memory. More work could be done to make this method run more efficiently. In addition, the performance of SIB ComBat may be sensitive to the choice of clusters per iteration as well as the number of iterations. For our real data analysis, we were able to leverage the MuSIC atlases to determine optimal cluster assignments based on relevant spatial information. However, additional work could be conducted to develop a streamlined way to determine the ideal cluster assignments and number of iterations if such information is not available. We also saw from the VBM analysis that a subset of features that were deemed statistically significant with respect to age in the original data were not statistically significant and vice-versa after performing harmonization with SIB-ComBat. Although the majority of associations were maintained, further work would aim to address this shortcoming (see Appendix for additional VBM analysis and simulation results).

A potential future direction would expand upon this framework to account for non-linear relationships with covariates. One way to do so would be to incorporate a generalized additive model, similar to ComBat-GAM [Pomponio et al., 2020]. Another possible future direction would look at the effectiveness of this method with other types of neuroimaging data. Here, we mainly were interested in the harmonization of high-dimensional, voxel-level intensity data, but it may be interesting to see how generally applicable this framework is for harmonizing various types of datasets, including longitudinal neuroimaging data. Furthermore, CAM ComBat could be expanded to include an additional prior on the batch specific covariance matrices. Because of the extra computational demand of introducing this prior, particularly in high dimensions, this direction was not explored in this paper. However, finding a way to introduce this prior in a computationally efficient way could be beneficial. In conclusion, CAM ComBat has shown to be an effective harmonization method to account for notable spatial heterogeneity between batches. In situations where you are working with hundreds of thousands of features, the extension of this method, SIB ComBat, is a promising alternative that both mitigates site-specific effects and maintains at least the same strength of biological associations found in the original data.

## Acknowledgements

The iSTAGING study is a multi-institutional effort funded by the National Institute on Aging (NIA). Data were provided by the Open Access Series of Imaging Studies (OASIS) and the Harvard Aging Brain Study (HABS).

## Appendix

### 5.1 Supplemental Calculations and Details

We first assume that *Y*_*ijv*_ can be modeled in the following way:

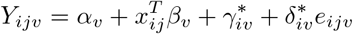

in which 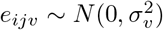. We first calculate the standardized data *Z*_*ijv*_ as

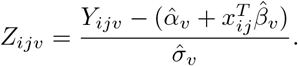

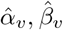 and 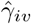are calculated using their respective voxel-wise ordinary least squares estimates. 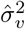 is calculated as 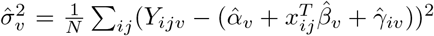, with *N* = Σ_*i*_*n*_*i*_.

Set *Z*_*ij*_ = (*Z*_*ij*1_, …, *Z*_*ijV*_). We will assume *Z*_*ij*_ ∼ *MV N* (*γ*_*i*_, Σ_*i*_) and introduce a multivariate normal prior such that 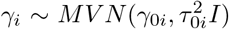. We use the method of moments estimators to estimate *γ*_*i*_, Σ_*i*_, *γ*_0*i*_, and 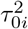 such that

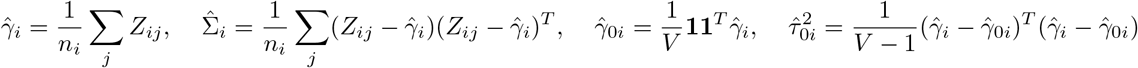

We can derive the conditional (posterior) distribution of *γ*_*i*_: *π*(*γ*_*i*_|*Z*_*i*_, Σ_*i*_)

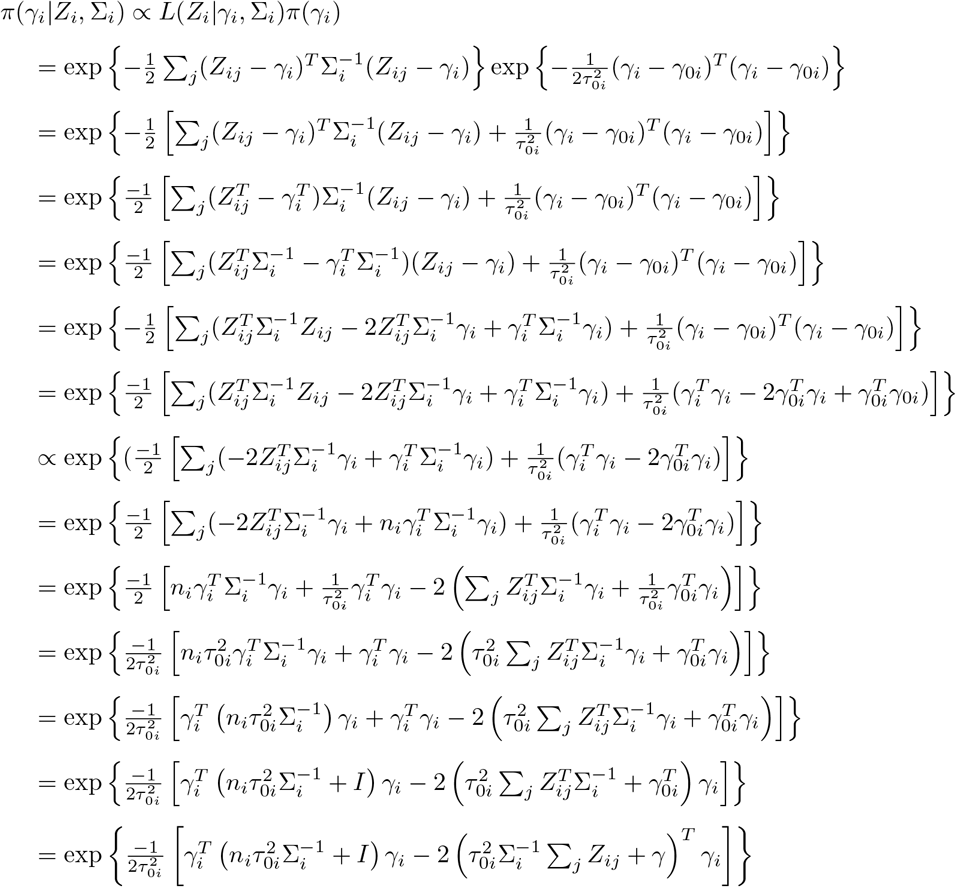

By completing the square, we can conclude that

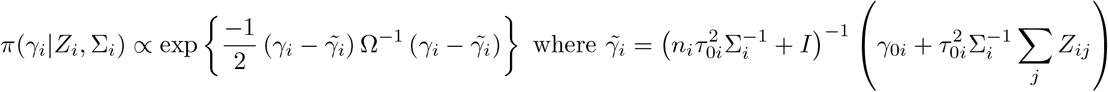

with matrix Ω calculated accordingly. This shows the posterior distribution is a multivariate normal distribution. We can then estimate *γ*_*i*_ based on the posterior mean, resulting in the following estimate:

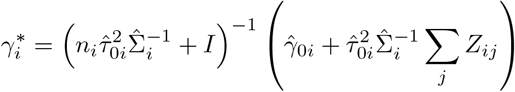

To model the batch-specific covariance effects, we first estimate the overall pooled covariance based on the sample covariance estimate: 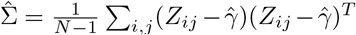, where *N* = Σ_*i*_*n*_*i*_ and 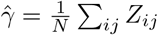. We then assume that 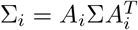, where *A*_*i*_ is a matrix dictating the batch-specific rotation. Leveraging the eigendecomposition of real symmetric matrices, we can rewrite the covariance matrices as 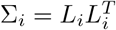 and Σ = *PP*^*T*^, in which

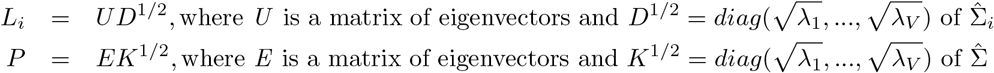

with *λ*_1_, …, *λ*_*V*_ as the eigenvalues of the respective covariance matrix. We can then write that 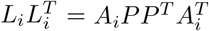 and subsequently *A*_*i*_ = *L*_*i*_*P* ^*−*1^. We finally adjust the data as follows to get the harmonized vector of imaging features *M*_*ij*_

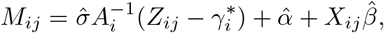

where 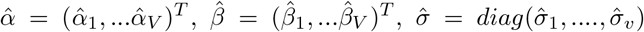, and *X*_*ij*_ is the matrix of covariate information.

### 5.2 Supplemental Simulation Results: Medium Difference

### 5.3 Supplemental Figure: The Effect of Different Harmonization Methods on Age Association

### 5.4 Supplemental VBM Analysis

### 5.5 Supplemental Figure: Comparison of Principal Component Loadings Between Sites

